# Sex classification from functional brain connectivity: Generalization to multiple datasets

**DOI:** 10.1101/2023.08.30.555495

**Authors:** Lisa Wiersch, Patrick Friedrich, Sami Hamdan, Vera Komeyer, Felix Hoffstaedter, Kaustubh R. Patil, Simon B. Eickhoff, Susanne Weis

## Abstract

Machine learning (ML) approaches are increasingly being applied to neuroimaging data. Studies in neuroscience typically have to rely on a limited set of training data which may impair the generalizability of ML models. However, it is still unclear which kind of training sample is best suited to optimize generalization performance. In the present study, we systematically investigated the generalization performance of sex classification models trained on the parcelwise connectivity profile of either single samples or a compound sample containing data from four different datasets. Generalization performance was quantified in terms of mean across-sample classification accuracy and spatial consistency of accurately classifying parcels. Our results indicate that generalization performance of pwCs trained on single dataset samples is dependent on the specific test samples. Certain datasets seem to “match” in the sense that classifiers trained on a sample from one dataset achieved a high accuracy when tested on the respected other one and vice versa. The pwC trained on the compound sample demonstrated overall highest generalization performance for all test samples, including one derived from a dataset not included in building the training samples. Thus, our results indicate that a big and heterogenous training sample comprising data of multiple datasets is best suited to achieve generalizable results.

## Introduction

Machine Learning (ML) is a powerful tool to relate neuroimaging data to behavior and phenotypes (Genon et al., 2022; Varoquaux & Thirion 2014) and is therefore increasingly being employed in neuroscience applications (Jollans et al., 2019; Buch et al., 2018; Varoquaux et al., 2018; Kohoutova et al. 2020). Successful applications of ML approaches include the decoding of mental states (Haynes & Rees 2006), classification of mental disorders (Zhang et al. 2021; Chen et al., 2020), as well as the prediction of demographic and behavioral phenotypes (Smith et al. 2015; Nostro et al., 2018; Pläschke et al., 2020; Varikuti et al., 2018; More et al., 2023).

ML models learn the feature-target relationship given a training sample. Subsequently, the model is applied to make predictions on previously unseen data (Dhamala et al., 2023) and successful generalization to independent data samples is the central goal in ML (Domingos, 2012; Varoquaux, 2018; Chung, 2018). For example, a recent study (Weis et al., 2020) demonstrated successful generalization of sex prediction models based on regionally specific functional brain connectivity patterns, which were trained on the data of the Human Connectome Project (HCP, Van Essen et al., 2012, Van Essen, 2013). For this spatially specific approach, independent classifiers were trained on the functional brain connectivity patterns of parcels covering the whole brain. In this case, assessing generalization performance should not only consider the averaged across-sample accuracy. Rather, if the classifiers generalize well, the same parcels should achieve high classification accuracies during cross-validation (CV) and across-sample testing.

Further sex classification studies (Menon & Krishnamurthy, 2019; Zhang et al., 2018; Smith et al., 2013), as well as other applications of ML models employed the HCP dataset to predict phenotypes such as task activation (Cohen et al., 2020), and individual behavioral and demographic scores (Smith et al., 2015; Cui & Gong, 2018) like age (Sanford et al., 2022). The HCP dataset is characterized by high-quality multi-modal imaging data acquired from a large group of healthy young adults. However, both the high quality of the brain imaging data as well as the narrow age range is not typical of other datasets, especially when dealing with clinical data (Arslan, 2018, Jansma et al., 2020; Rutten et al., 2010). This raises the question whether results based on the HCP data can be generalized to other datasets with different characteristics. Weis et al., (2020) demonstrated that sex classifiers trained on the HCP data generalized well to an independent subset of the HCP dataset as well as to the 1000Brains dataset (Caspers et al. 2014). Additional evidence from the application of such classifiers to data from datasets with diverse characteristics would provide even stronger evidence of model generalization.

Especially in neuroimaging, differences between datasets may result from several different sources. On the one hand, participants may differ with respect to demographic characteristics, such as age, education, or economic status. On the other hand, data samples likely differ with regard to the MRI acquisition parameters and data processing. Considering these differences, it is so far unresolved what kind of training sample leads to good generalization performance across multiple test samples.

Various characteristics of the training data can influence the generalization performance of ML models (Dhamala et al. 2023). For instance, larger sample size is beneficial for generalization performance (Cui & Gong, 2018, Domingos, 2012). Ensuring that the training data is representative of the target sample is another crucial factor for achieving good generalization performance (Ishida, 2019, Yang et al. 2020). Furthermore, data from different acquisition sites are likely heterogeneous with respect to demographic characteristics, data acquisition, and processing parameters. Therefore, a ML model trained on such data is less likely to overfit. Thus, aggregating data from multiple sites should be beneficial for improving generalization performance. Indeed, this has been partially shown by studies concerning clinical applications of ML approaches (Nielsen et al, 2020; Willemink et al. 2020; Chang et al. 2018). These results suggest that training ML models on diverse datasets covering a wide range of characteristics may improve the overall generalization performance.

In the present study, our aim was to evaluate the generalization performance of sex classifiers trained on samples created from four different datasets with varying demographic characteristics. In addition, sex classifiers were trained on a compound sample combining data from all datasets to obtain a training sample with heterogeneous sample characteristics. Following the parcelwise approach by Weis et al. (2020), we trained independent sex classifiers on the resting state (RS) connectivity patterns of 436 parcels covering the whole brain. For each parcel, a sex classification model was built based on the individual connectivity profile, resulting in one classification accuracy value per parcel. This was done for each of the five training samples, resulting in five sets of parcelwise classifiers (pwCs). These pwCs were applied to test samples from the four original datasets and one dataset which was not part of the training samples. Then, accuracy maps, representing the spatial distribution of classification accuracies for each parcel were generated for CV (within-sample accuracy) and for application of the pwCs to the different test samples (across-sample accuracy). The comparison of these accuracy maps enabled us to evaluate generalization performance of classifiers by (i) examining the mean accuracy of all parcelwise classifiers across the 10% best classifying parcels and (ii) comparing the spatial location of highly classifying parcels between CV and across-sample test. Good generalization performance with regard to spatial consistency is characterized by identical parcels performing well in CV and across-sample testing. We hypothesized that the compound sample achieves best generalization performance as suggested by previous literature (Nielsen et al, 2020; Willemink et al. 2020; Chang et al. 2018).

## Materials and Methods

### Data

We employed resting state functional magnetic resonance imaging (fMRI) data of subsets of four large datasets to train and test sex classification models. For all datasets, we only included healthy subjects aged 20 years or older. Within each training sample, we matched females and males for age and included a similar number of women and men. The first sample, taken from the HCP dataset (900 subjects data release; Van Essen et al., 2012; Van Essen 2013), comprised 878 subjects with a mean age of 28.49 years (range: 22-37 years). The second sample, taken from the Brain Genomics Superstruct Project (GSP; Holmes et al., 2015) comprised 854 subjects with a mean age of 22.92 years (range: 21 – 35 years). The third sample was a subset from the Rockland Sample of the Enhanced Nathan Klein Institute (eNKI; Nooner et al., 2012), comprising 190 subjects with a mean age of 46.02 years (range: 20-83 years). The fourth sample, taken from the 1000Brains dataset (Caspers et al., 2014), comprised 1000 subjects with a mean age of 61.18 years (range: 21-85 years). This sample was included to examine generalization performance to an older sample. A fifth sample (“compound”) was constructed by combining 75% of the HCP, GSP, eNKI and 1000Brains samples. The compound sample comprised an age range of 20-85 years (*M* = 40.10, *SD* = 19.96 years). RS fMRI data from an additional dataset was included to evaluate classifiers on an additional independent sample. This sample comprised 370 subjects (214 females) with a mean age of 22.50 years (range 20-26 years) from the AOMIC dataset (Snoek et al., 2021). It was not additionally balanced for sex to maintain the maximum number of participants for evaluation. Data usage of the included datasets was approved by the Ethics Committee of the Medical Faculty of the Heinrich-Heine University Düsseldorf (4039, 5193, 2018-317-RetroDEuA). All data was collected in research projects approved by a local Review Board, for which all participants provided written informed consent. All experiments were performed in accordance with relevant guidelines and regulations.

### Data acquisition and preprocessing

#### HCP

The RS fMRI data of the HCP dataset were acquired on a Siemens Skyra 3T MR scanner with multiband echo-planar imaging with a duration of 873 seconds and the following parameters: 72 slices; voxel size, 2 x 2 x 2 mm^3^; field of view (FOV), 208 x 180 mm^2^; matrix, 104 x 90; TR,

720 ms; TE, 33 ms; flip angle, 52 degrees (https://www.humanconnectome.org/storage/app/media/documentation/s1200/HCP_S1200_Release_Reference_Manual.pdf) Participants were instructed to lie in the scanner with eyes open, with a “relaxed” fixation on a white cross on a dark background and think of nothing in particular, and to not fall asleep (Smith et al., 2013). RS data were corrected for spatial distortions, head motion, B_0_ distortions and were registered to the T1-weighted structural image (Smith et al. 2013). Concatenating these transformations with the structural-to-MNI nonlinear warp field resulted in a single warp per time point, which was applied to the timeseries to achieve a single resampling in the 2mm MNI space. Afterwards, global intensity normalization was applied and voxels that were not part of the brain were masked out. Locally noisy voxels as measured by the coefficient of variation were excluded and all the data were regularized with 2mm Full width half maximum (FWHM) surface smoothing (Smith et al. 2013; Glasser et al. 2013). The temporal preprocessing included corrections and removal of physiological and movement artifacts by an independent component analysis (ICA) of the FMRIB’
ss X-noisifier (FIX, Salimi-Khorshidi et al., 2014). This method decomposes data into independent components and identifies noise components based on a variety of spatial and temporal features through pattern classification.

#### GSP

RS data were acquired on a 3T Tim Trio Scanner with a duration of 372 seconds and the following parameters: 47 slices; voxel size, 3 x 3 x 3 mm^3^; FOV read, 216 mm; TR, 3 s; TE, 30 ms; flip angle, 85 degrees. During data acquisition, participants were instructed to lay still, stay awake, and keep eyes open while blinking normally (https://static1.squarespace.com/static/5b58b6da7106992fb15f7d50/t/5b68650d8a922db3bb807a90/1533568270847/GSP_README_140630.pdf, Holmes et al. 2015).

#### eNKI

Participants in the eNKI dataset were underwent RS scanning for 650 seconds in a Siemens Magnetom Trio Tim sygno MR scanner with the following parameters: 38 slices; voxel size, 3 x 3 x 3 mm^3^, FOV, 256 x 200mm^2^; TR, 2500 ms; TE, 30 ms; flip angle, 80 degree. Participants were instructed to keep their eyes closed, relax their minds and not to move (Betzel et al. 2014).

#### 1000Brains

Subjects were scanned for 660 seconds on a Siemens TRIO 3T MRI scanner with the following parameters: 36 slices; voxel size, 3.1 x 3.1 x 3.1 mm^3^; FOV, 200 x 200 mm^2^; matrix, 64 x 64, TR = 2.2 s; TE = 30 ms; flip angle, 90 degrees. During RS data acquisition, participants were instructed to keep their eyes closed and let the mind wander without thinking of anything in particular (Caspers et al. 2014).

RS data of the GSP, eNKI and 1000Brains samples were preprocessed in the same way. The preprocessing pipeline comprised removal of noise and motion artifacts by using FIX (Salimi-Khorshidi et al., 2014). The denoised data were further preprocessed with SPM12 (SPM12 v6685, Wellcome Centre for Human Neuroimaging, 2018; https://www.fil.ion.ucl.ac.uk/spm/software/spm12/) using Matlab R2014a (Mathworks, Natick, MA). For each subject, the first four echo-planar-imaging (EPI) volumes were discarded and the remaining ones were corrected for head movement by an affine registration with two steps: First, the images were aligned to the first image. Second, the images were aligned to the mean of all volumes. The resulting mean EPI image was spatially normalized using the MNI152 template (Holmes et al., 1998) using the “unified segmentation” approach in order to take into account inter-individual differences in brain morphology (Ashburner & Friston, 2005). After deformation application and smoothing, images were resampled to partial volume label image to 2x2x2mm^3^ and resampled using the modulated GM segment image to 2x2x2mm^3^.

#### AOMIC

The AOMIC dataset includes two subsamples, PIOP1 and PIOP2, comprising data of healthy university students scanned on a Philips 3T scanner. Participants were instructed to keep their gaze fixated on a fixation cross on the screen and let their thoughts run freely (Snoek et al., 2021). Both samples were acquired with a voxel size of 3 x 3 x 3 mm^3^ and a matrix size of 80 x 80. While PIOP1 was acquired for 360 seconds with multi-slice acceleration, 480 volumes and a 0.75 TR, PIOP2 was acquired for 480 seconds without multi-slice acceleration, 240 volumes and a 2s TR (further details in https://www.nature.com/articles/s41597-021-00870-6/tables/10). Data were preprocessed using Fmriprep version 1.4.1 (Esteban et al., 2019; Esteban et al., 2020), a Nipype based tool for reproducible preprocessing in neuroimaging data (Gorgolewski et al., 2011). Data were motion corrected using mcflirt (FSLv5.0.9, (Jenkinson et al. 2002)) followed by distortion correction by co-registering the functional image to the respective T1 weighted image with inverted intensity (Huntenburg, 2014; Wang et al., 2017) with 6 degrees of freedom, using bbregister (FreeSurfer v6.0.1). In a following step, motion correction transformations, field distortion correction warp, BOLD-to-T1-weighted transformation and the warp from T1-weighted to MNI were concatenated and applied using antsApplyTransforms (ANTs v2.1.0.) using Lanczos interpolation (Snoek et al., 2021).

### Connectome extraction

Following the parcelwise approach by Weis et al. (2020), individual RS connectomes were extracted based on 400 cortical parcels of the Schaefer Atlas (Schaefer et al. 2018), and 36 subcortical parcels of the Brainnetome Atlas (Fan et al., 2016). Each parcel’s time series was cleaned by excluding variance that could be explained by mean white matter and cerebrospinal fluid signal (Satterthwaite et al., 2013). Data was not further cleaned for motion related variance as this variance was already removed during FIX preprocessing. For each of the 436 parcels, the activation time series was computed as the mean of all voxel time courses within that parcel. Then, for each parcel, pairwise Pearson correlations were computed between the parcel’s time series and those of all other 435 remaining parcels, representing the individual RS functional connectivity (RSFC) profile of the parcel.

### Parcelwise sex classification

Sex classification models were trained based on the individual multivariate RSFC profile of each parcel, resulting in a set of 436 pwC (Weis et al., 2020). All models were built using support vector machine (SVM) classifiers. SVM is a supervised ML method that separates the data into distinct classes – males and females in case of sex classification – with the widest possible gap between these classes (Vapnik 1998; Boser et al., 1992; Rafi & Shaikh., 2013; Zhang et al., 2021). SVM models were built in Julearn (https://juaml.github.io/julearn/main/index.html) including a Hyperparameter search nested within a 10–fold CV with 5 repetitions. The parameter search included choice of kernel (linear vs. radial basis function (rbf) kernel) as well as the C- and gamma-parameter. Confounding effects of age were regressed out by removing age-related variance before training the classifiers. The best performing combination of hyperparameters was used for the final model for each individual parcel. Within-sample classification accuracy for each individual parcel was determined by averaging accuracies over CV folds and repetitions.

For across-sample classification, single dataset pwCs were tested on the respective other three samples (sample characteristics displayed in table 1, Figure 1), while the compound sample pwC were tested on the remaining 25% of the HCP (n = 220, age range: 22-36), GSP (n = 214, age range: 21-31), eNKI (n = 48, age range: 20-75) and 1000Brains (n = 250, age range: 22-80) sample (Table 1). Here, for computing time reasons, we restricted the choice of the SVM kernel to rbf (see Weis et al., 2020). Finally, generalization performance of all five pwCs was assessed on the AOMIC sample. All reported accuracies are balanced accuracies.

**Table 1.**
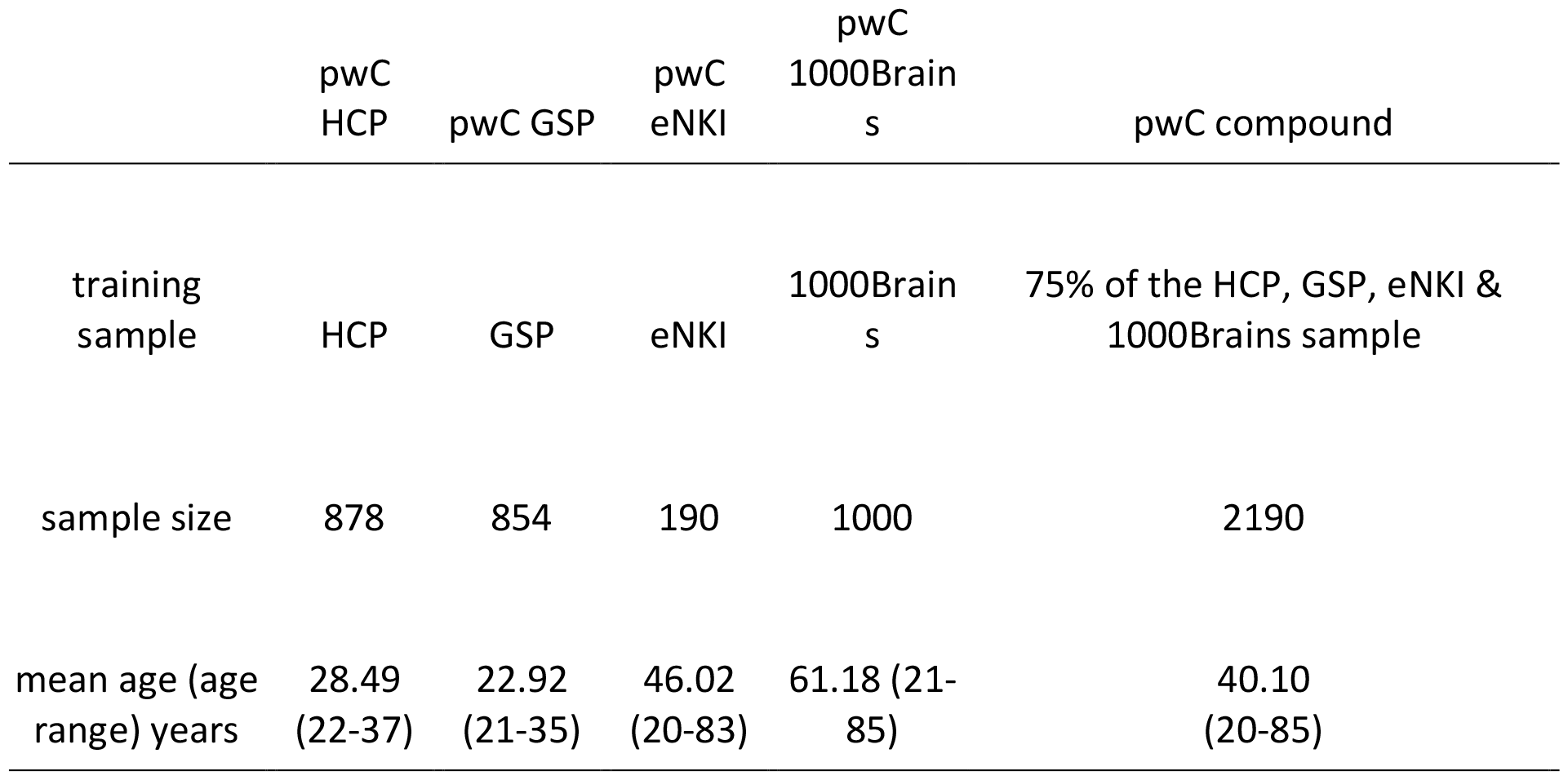
Sample characteristics. Depiction of sample characteristics of each sample to train the respective pwC.

|  | pwC<br>HCP | pwC GSP | pwC<br>eNKI | pwC<br>1000Brain<br>s | pwC compound |
| --- | --- | --- | --- | --- | --- |
| training<br>sample | HCP | GSP | eNKI | 1000Brain<br>s | 75% of the HCP, GSP, eNKI &<br>1000Brains sample |
| sample size | 878 | 854 | 190 | 1000 | 2190 |
| mean age (age<br>range) years | 28.49<br>(22-37) | 22.92<br>(21-35) | 46.02<br>(20-83) | 61.18 (21-85) | 40.10<br>(20-85) |

**Figure 1.**
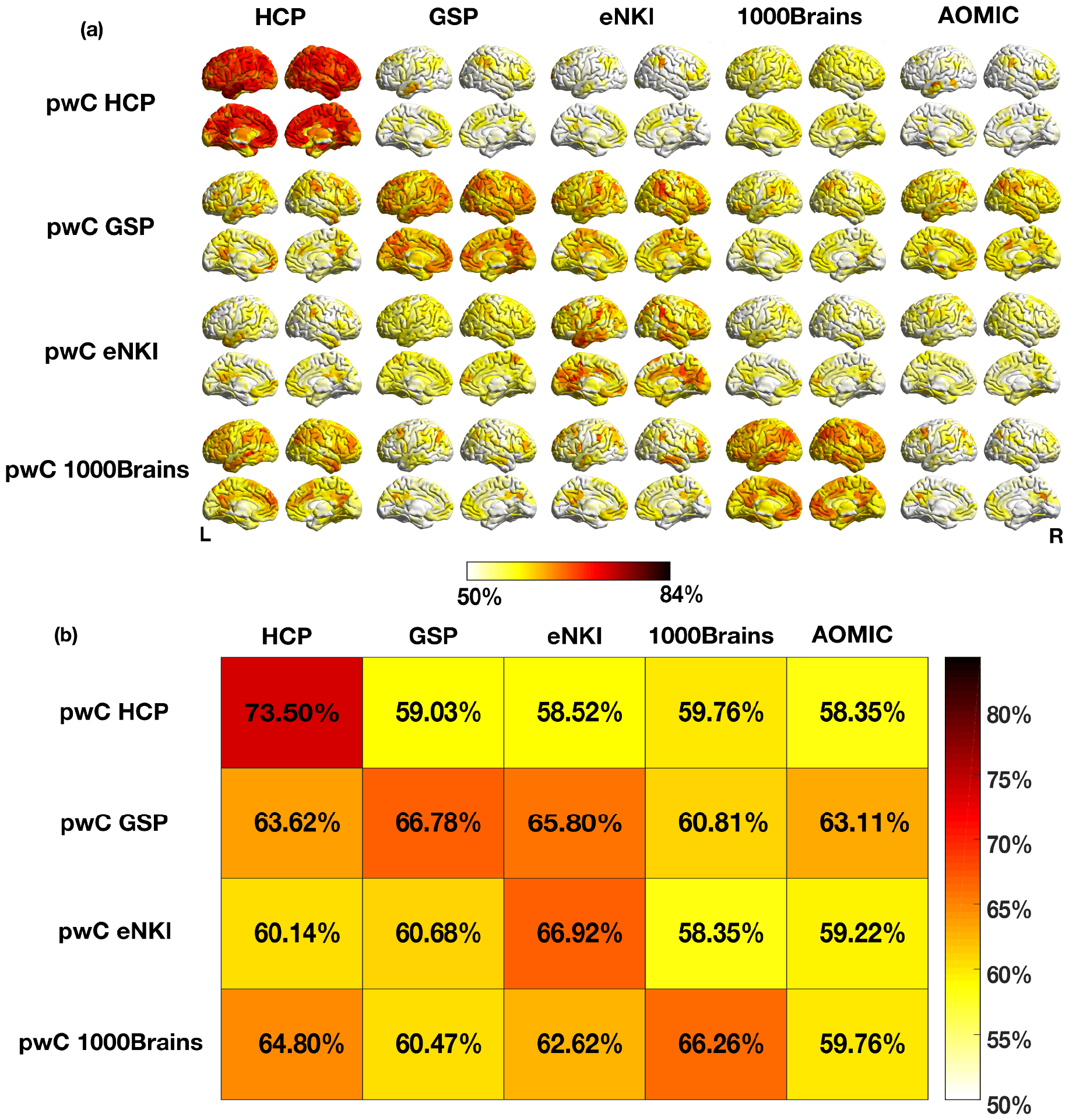
Accuracy maps and tile plots of mean accuracies of top 10% classifying parcels for pwCs trained on single samples. (a) Spatial distribution of parcelwise sex classification accuracies across the brain. Within-sample accuracies are depicted on and across-sample accuracies off the diagonal. Only parcels with an accuracy of 0.5 or higher are displayed. (b) Mean accuracies averaged across the top 10% classifying parcels for each CV- and across-sample prediction.

## Statistical analyses

### Across-sample classification accuracy

To statistically compare the classification accuracies achieved by each pwC on the different test samples, we employed independent t-tests between the different across-sample accuracies over all 436 parcels. To compare the performance of the different pwCs on each test sample, independent t-tests across all parcels were used. Significance levels were Bonferroni-corrected according to the number of dependent tests (10 dependent tests for comparing the across-sample accuracy of pwC compound on the five fest samples as well as for comparing the across-sample accuracies of the five pwCs on the AOMIC sample; 6 dependent tests for all other comparisons).

### Consistency of highly classifying brain regions

Previous studies have demonstrated that sex classification accuracies for models trained on parcelwise RSFC patterns do not achieve uniformly high performance across the whole brain (Weis et al. 2020; Zhang et al. 2018). Thus, we assessed generalization performance of the different pwCs by examining the consistency of highly classifying brain regions during CV and across-sample testing. Consistency was assessed by computing Dice coefficients (DSC) to evaluate the similarity in spatial distribution of parcels achieving certain accuracies in both CV and across-sample testing. This consistency was evaluated for different accuracy thresholds above chance (0.5 - 0.7 at 0.02 steps). For each threshold, Dice coefficients were computed as the number of common parcels achieving within- and across-sample accuracies above or equal to that threshold (p_com) multiplied by 2 and divided by the total number of parcels achieving a within (p_tr) or across-sample (p_te) accuracy above or equal to that accuracy level in CV (Dice, 1945; Sorensen, 1948).

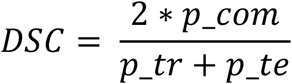

To facilitate comparison of the dice score distributions between the different pwCs and test samples, we summarized each contribution into one score by computing a weighted mean (wmDice) as the average of each dice coefficient weighted by the accuracy threshold for which the respective dice coefficient was calculated.

## Results

The generalization performance of pwCs trained on each of the single dataset samples (HCP, GSP, eNKI, & 1000Brains) and on the compound sample were compared with respect to mean across-sample accuracy averaged across the best 10% classifying parcels. Additionally, we evaluated the consistency of the spatial distribution of accurately classifying parcels between CV and across-sample testing.

### Training and test classification accuracies

For the single samples pwCs, the mean within-sample performance across the top 10% classifying parcels was at a similar level for pwC GSP (66.8%), pwC eNKI (66.9%) and pwC 1000Brains (66.3%) and ranged up to 73.5% for pwC HCP. The mean across-sample accuracies averaged for the top 10% classifying parcels ranged between 58.4% (for pwC HCP tested on AOMIC and pwC eNKI tested on 1000Brains) and 65.8% (for pwC GSP tested on eNKI). Details for within- and across-sample performance are reported in Table S1 and Figure 1 and Figure S1. Parcelwise within- and across-sample accuracies are displayed as accuracy maps in figure 1a and the distribution of test accuracies is shown in figure 2 (red boxplots). Here, accuracy maps represent the spatial distribution of classification accuracies resulting from the 436 individual ML models trained on the respective multivariate RSFC profile of each parcel.

**Figure 2.**
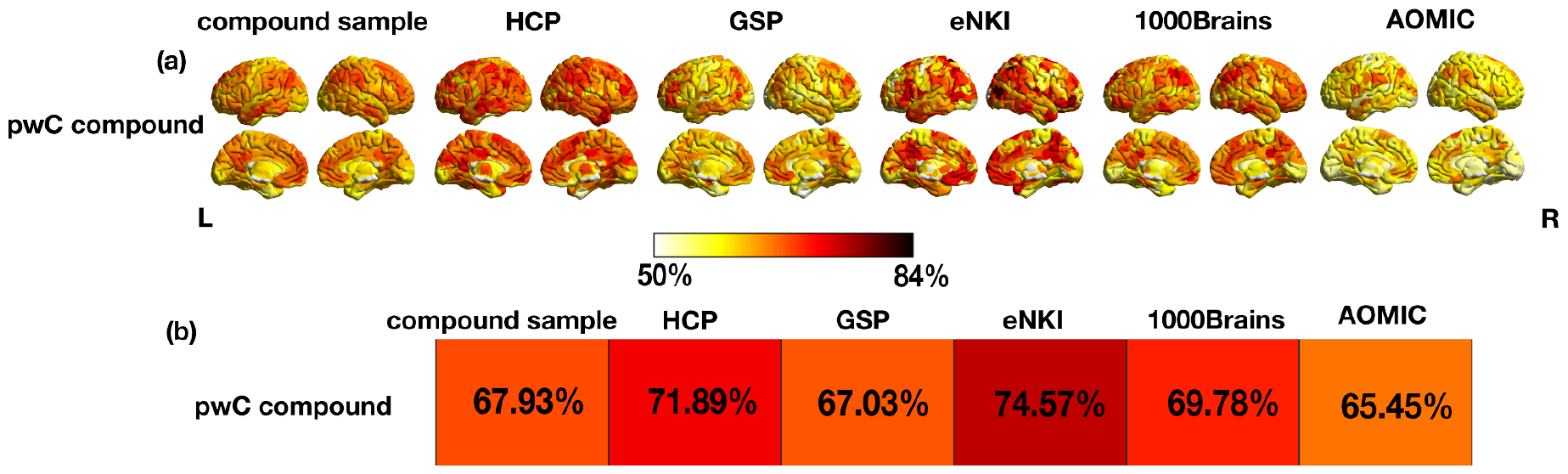
Accuracy maps and tile plots of mean accuracies of top 10% classifying parcels for pwC compound. (a) Spatial distribution of parcelwise sex classification accuracies across the brain. Only parcels with an accuracy of 0.5 or higher are displayed. (b) Mean accuracies averaged across the top 10% classifying parcels for the CV- and across-sample prediction.

Accuracy maps for the different combinations of training and test samples were compared using independent t-tests across the top 10% classifying parcels in each prediction (details in Table S2). First, we analyzed differences in classification accuracies between test samples for each pwC (horizontal comparisons, figure 1): For pwC HCP, testing on 1000Brains achieved the highest mean classification accuracy (59.8%). The averaged accuracy for this test sample was descriptively higher than for the GSP and significantly higher than for the eNKI and AOMIC test samples. PwC GSP achieved significantly higher accuracies for the eNKI test sample (65.8%) than for any other test sample, while pwC eNKI showed highest accuracies for the GSP test sample (60.7%). This across-sample prediction showed descriptively higher accuracies than pwC eNKI did for the HCP test sample and significantly higher accuracies than for the AOMIC and 1000Brains samples. For pwC 1000Brains, testing on the HCP showed significantly higher accuracies (64.8%) than testing on the eNKI, GSP and AOMIC sample. Details of all statistical comparisons are given in Table S2.

PwC compound achieved a mean within-sample accuracy of 67.9% within the top 10% classifying parcels. The mean across-sample accuracies averaged across the top 10% classifying parcels ranged between 65.5% (pwC compound tested on AOMIC) and 74.6% (pwC compound tested on eNKI, details in Table S1 and Figure 2, Figure S2).

Contrasting the top 10% classifying parcels in the accuracy maps of pwC compound displayed peaks in accuracies for the eNKI test sample (74.6%) resulting in significantly higher accuracies than for the remaining test samples (Figure 2 and Table S2). We also contrasted how the five pwCs performed on each test sample by employing independent t-tests: pwC compound outperformed all pwCs trained on single dataset samples for the HCP, GSP, eNKI and AOMIC test sample with regards to the top 10% classifying parcels in each across-sample prediction. Details for all statistical comparisons are shown in Table S2.

### Consistency of correctly classifying parcels

**Figure 2.**
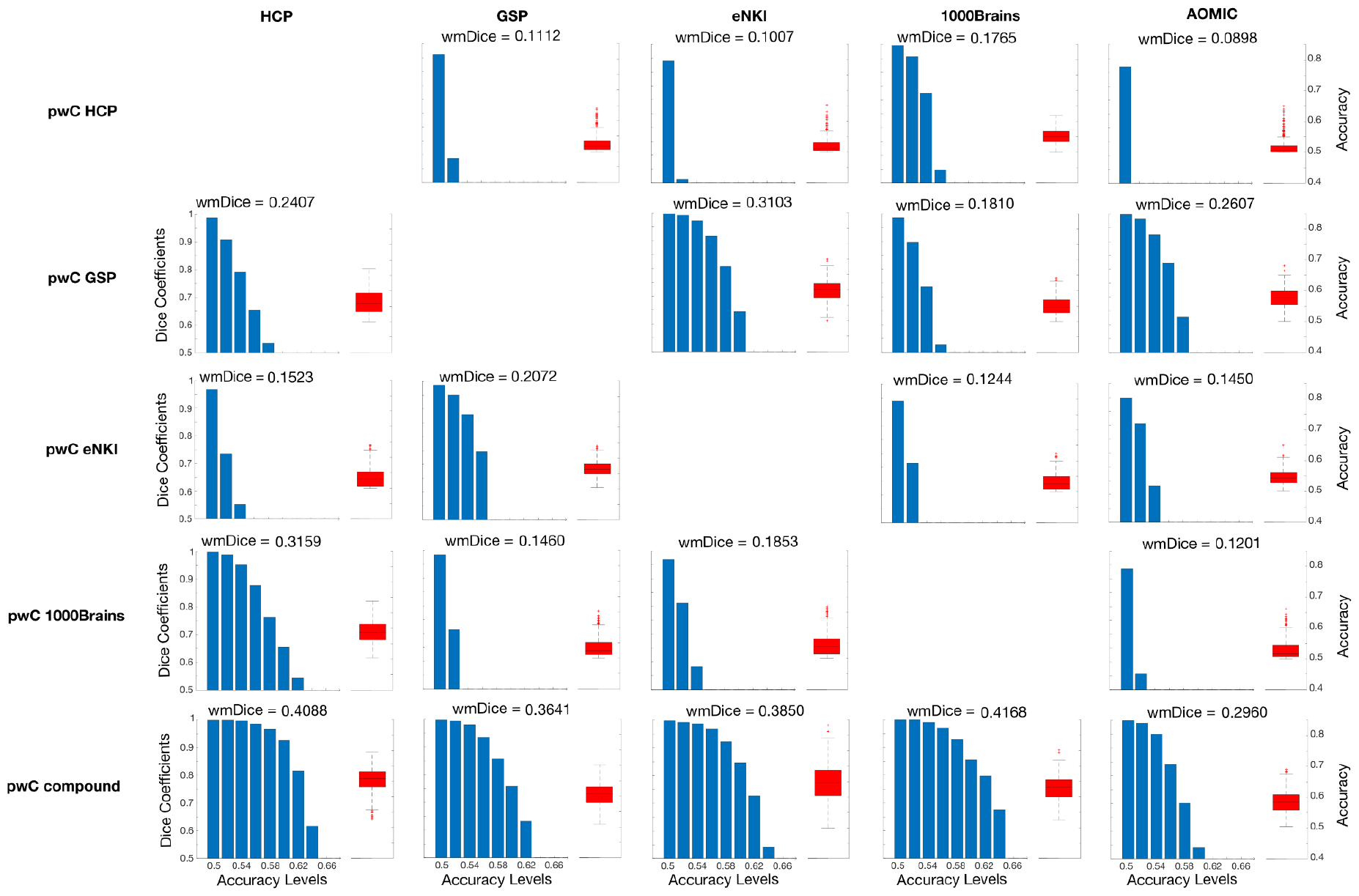
Spatial consistency of all pwCs. For each combination of training (rows) and test sample (columns), the right side of each subplot (red boxplot) depicts the distribution of accuracies across all parcels (right y-axis). The left side of each subplot (blue barplot) shows the dice coefficients (left y-axis), representing the overlap of accuracy maps between CV and test at different accuracy levels (x-axis). For each accuracy-threshold, the respective dice coefficient was calculated as the number of similar parcels classifying above a certain accuracy-threshold in both, respective CV and test prediction, in relation to the total number of parcels of both predictions classifying at this level. For each combination of pwC and test sample, the weighted mean of the dice coefficients (wmDice) across accuracy levels is displayed above the subplot to allow for a straightforward comparison between the distributions of dice coefficients.

To evaluate the spatial consistency of accurately classifying parcels, we calculated the dice coefficient between thresholded within- and across-sample accuracy maps at different levels of accuracy. Here, a high dice coefficient indicates a high overlap in highly classifying parcels between within and across-sample predictions at a given accuracy level. The results are depicted in the blue bar plots in figure 2. Regarding spatial consistency within a given pwC (horizontal comparison in Fig 2), pwC HCP overall demonstrated relatively low spatial consistency while it was highest for 1000Brains (wmDice = 0.1765, all other wmDice < 0.1112). Spatial consistency for pwC GSP was highest for the eNKI sample (wm = 0.3103) and lowest for 1000Brains (wmDice = 0.1810) with spatial consistency for HCP (wmDice = 0.2407) and AOMIC (wmDice = 0.2607) test samples ranging in between. PwC eNKI showed overall low spatial consistency for the HCP, 1000Brains and AOMIC sample (wmDice: 0.1244 - 0.1523) and highest for the GSP sample (wmDice = 0.2072). Spatial consistency of pwC 1000Brains was lower for the GSP, eNKI and AOMIC test sample (wmDice: 0.1201 - 0.1853) but considerably higher for the HCP test sample (wmDice = 0.3159) pwC compound demonstrated relatively similar spatial consistency for HCP, GSP, eNKI and 1000Brains (wmDice: 0.3641 - 0.4168) and lower spatial consistency with the AOMIC sample (wmDice: 0.2960). Concerning the comparisons within each test sample (vertical comparisons in Fig 2) pwC compound demonstrated higher spatial consistency than all single dataset sample pwCs for all test samples. Dice coefficients for the top 10% classifying parcels are reported in figure S3.

## Discussion

In the present study, we examined the generalization performance of parcelwise sex classification models trained on different samples. Here, we operationalized generalization performance in terms of both mean classification accuracy of best classifying parcels during across-sample testing as well as spatial consistency in highly classifying parcels between CV and across-sample test. Since not all parcels can be expected to achieve high classification accuracies, we mainly focused on the top 10% classifying parcels. Overall, our results showed that classifiers trained on single dataset samples generalized well only for certain, but not for all, test samples. In contrast, classifiers trained on the compound sample outperformed classifiers trained on single dataset samples both in terms of accuracy and consistency of accurately classifying parcels.

To evaluate generalization performance with respect to mean classification accuracies of the top 10% classifying parcels, for each pwC, we compared across-sample classification accuracies between the different test samples. Results indicate that certain datasets seem to “match” in the sense that classifiers trained on a sample from one of the datasets achieved a high accuracy when tested on the respective other one and vice versa. This was the case for HCP and 1000Brains as well as for GSP and eNKI with the former matching the results of a previous study (Weis et al., 2020). Based on the good across-sample performance of sex classifiers trained on an HCP sample on a subsample of the 1000Brains, Weis et al., (2020) suggested that parcelwise sex classification generalizes well between different samples. No additional samples from other datasets were considered in Weis et al. (2020). The present results extend the findings of the previous study by showing that good generalization performance of the HCP classifiers appears to be specific to the 1000 Brains sample. Generalization to samples from other datasets (GSP, eNKI and AOMIC) is, however, rather poor. Thus, our study demonstrates that the generalizability of pwCs trained on single dataset samples depends on the train-test data combination, which is in line with a previous study that employed sex classification based on regional homogeneity of resting state time series (Huf et al., 2014). The limited generalization performance of the pwCs trained on single dataset samples to the majority of test samples from other datasets might be attributed to the homogeneity of each single dataset training sample arising due to demographic factors such as the age range (Damoiseaux et al., 2008; Damoiseaux, 2017; Scheinost et al., 2015) as well as technical details such as fMRI acquisition parameters (Yu et al., 2018; Brown et al., 2011). Homogeneous data characteristics within each dataset will result in a homogeneity of the feature space on which ML models are trained. Such homogeneous features might lead the ML model to learn dataset specific characteristics that are predictive of the target variable, which might not translate to other test samples, resulting in inaccurate across-sample predictions (Huf et al., 2014). Thus, training ML models on a single, homogenous sample may not be ideal to achieve a good generalization performance on diverse test samples (Huf et al., 2014; Janssen et al., 2018; Belur Nagaraj et al., 2020; Di Tanna et al., 2020). In contrast, training classifiers on a combination of multiple datasets achieved significantly higher accuracies for all test samples, including the sample from a dataset which was not included in the compound training sample. The increased generalization performance might be attributable to the heterogeneity of data characteristics included in a training sample created from various datasets. This heterogeneity likely enables the model to learn patterns that do not rely on specific sample characteristics, but actually capture the underlying relationship between features and target, enabling the model to generalize better, even to data from datasets that were not included in training. Thus, training on a compound sample is preferable to training on single dataset samples (Huf et al., 2014; Chang et al., 2018; Willemink et al., 2020).

The parcelwise classification approach allowed us to investigate generalization performance not only in terms of accuracy but also with respect to the spatial distribution of accurately classifying parcels. To quantify the overlap of accurately classifying parcels between CV and across-sample test, we computed dice coefficients between within- and across sample accuracy maps at different accuracy thresholds. We observed a pattern similar to the one found for classification accuracies, with the train-test pairing of HCP and 1000Brains and GSP and eNKI, respectively, showing highest spatial consistency, relative to other combinations. Thus, also when considering spatial consistency, generalization performance depended on specific pairing of training and test datasets. For pwCs trained on single samples, training sample characteristics appeared to be the most important factor in driving generalization performance across test samples. In contrast, pwC compound achieved superior spatial consistency across all test samples, as compared to pwCs trained on single samples. Thus, the classifiers trained on the compound sample achieved both higher classification accuracies as well as more consistency in accurately classifying parcels as opposed to the classifiers trained on single dataset samples. Altogether, the high generalization performance for pwC compound can likely be attributed to the heterogeneity in the compound sample which was achieved by combining multiple samples for training. These findings match results of previous studies (Huf et al., 2014; Chang et al., 2018; Nielsen et al., 2020; Willemink et al., 2020).

Overall, the aggregation of multiple samples in pwC compound for training sex classifiers resulted in superior generalization performance. Firstly, the classification accuracies were comparable between CV and the different across-sample test classifications. Secondly, highly classifying parcels overlapped to a large degree between training and and test. The overall high generalization performance of pwC compound across all test samples could be attributed to several possible explanations: first, the compound sample is more than twice as large as compared to any of the single dataset samples. Such high sample size has been shown to be beneficial for generalization (Cui & Gong, 2018, Domingos 2012, Ishida, 2019, Yang et al. 2020). However, sample size alone is likely not sufficient to explain the high generalization performance. For instance, the eNKI sample consists of only 190 participants, but the classifiers trained on this sample achieved better generalization performance than those trained on the HCP sample, which included 878 participants. A second explanation for the good performance of pwC compound may lie in the heterogenous nature of its training sample as discussed above. Having the different samples represented within the compound sample may have allowed the classifiers to classify sex based on sample-unspecific information. Another potential explanation is that the training sample of pwC compound partially consists of data from datasets on which we evaluated the test performance. In general, training on data that is representative of the test data typically results in an increased generalization performance (Chung et al., 2018). In contrast to pwC compound, CV and across sample test performances differed considerably for pwCs trained on single dataset samples. This lack of generalization performance was especially apparent for pwC HCP which showed a rather high performance during CV in combination with the lowest generalization performance both with respect to accuracy and spatial consistency. While homogeneity of a data sample has been argued to lead to high CV classification accuracy (Huf et al., 2014), sample characteristics such as the age range were comparable between HCP and the GSP sample, with the latter outperforming HCP in generalization performance. Thus, the comparably poor performance of classifiers trained on the HCP sample may be partially attributed to sample homogeneity but also to other factors such as the differences in preprocessing pipelines. For the HCP sample, connectome extraction was based on the minimally preprocessed version of the data. The eNKI, GSP and 1000Brains samples were preprocessed using the same pipeline in SPM12, while the AOMIC sample was preprocessed using fMRIprep. Given that comparative performance evaluation of fMRI data is sensitive to preprocessing decisions (Bhagwat et al., 2021), it is likely that this difference in preprocessing may contribute to the poor generalization performance of pwC HCP when tested on the other single samples. Furthermore, the high within-sample accuracy coupled with the lack of generalization performance may also indicate an overfitting effect of pwC HCP during training (Domingos, 2012; Cui & Gong, 2018). Altogether, our results highlight the importance of a heterogenous, diverse, and representative data composition for training ML models (Gong et al., 2019; Li et al., 2022; Dhamala et al., 2023), which can be achieved by combining data from multiple sites and datasets (Nielsen et al., 2020; Willemink et al., 2020; Chang et al., 2018). By minimizing sample-specific biases, we can aim for maximizing the generalizability of ML models.

### Limitations

The present results consistently demonstrated the superior generalizability of sex classifiers trained on a compound sample as compared to those trained on single dataset samples, but they come with some limitations. First of all, the high spatial consistency of pwC compound might partially be attributed to the generally higher accuracy of the across-sample predictions. Dice coefficients across the top 10% classifying parcels showed a more differentiated pattern. Here, pwC compound did not always outperform pwCs trained on single samples.

Another limitation in the present study is that, while we accounted for age as a potential confound during training of the classifiers, there might be other confounds that were not considered. For example, we did not control for structural variables such as brain size, which have been reported to influence brain functions (Batista-García-Ramó, K., & Fernández-Verdecia, C. I., 2018) and RS brain connectivity in particular (Zhang et al., 2018). Thus, in principle, different distributions of brain size within the different samples might have influenced the present results. However, Weis et al. (2020) demonstrated that at least with their training sample, classification based on RS connectivity was not systematically influenced by brain size. Still, there might be other demographic variables which differ between samples and might influence classification accuracies (Sripada et al., 2021; Mehrabi et al., 2021; Li et al., 2022).

Another factor which has not been considered in the present analyses are fluctuating sex hormones, which have been shown to influence functional brain connectivity in RS (Weis et al., 2019; Arélin et al. 2015; Haraguchi et al. 2021). These dynamic changes in female and male connectivity patterns (Ewen & Milner, 2017; Coenjaerts et al., 2023; Kogler et al., 2016) will likely influence overall sex classification accuracies. However, unfortunately, most publicly available datasets do not provide information on hormone levels, making it impossible to consider these variations in the analyses. Future large-scale studies should include hormone levels in data acquisition, enabling model training on a combination of multiple independent datasets with well characterized phenotypes to achieve most accurate results.

## Conclusion

The present results show that parcelwise sex classification models generalize best when trained on a compound sample including data with different demographic and data acquisition characteristics. Our results demonstrate that a large and heterogenous training sample including multiple datasets is best suited to achieve accurate generalization performance. This observation carries practical implications for future neuroimaging studies employing ML models for generalizable predictions.

## Acknowledgements

We would like to thank Dr. Federico Raimondo and Dr. Georgios Antonopoulos and all people contributing in the “ML hour” initiative from INM-7 for the insightful discussions.

## Data availability statement

The datasets HCP, GSP, eNKI and AOMIC are publicly available and free to download:

https://www.humanconnectome.org/study/hcp-young-adult/data-releases

https://dataverse.harvard.edu/dataset.xhtml?persistentId=doi:10.7910/DVN/25833

https://openneuro.org/datasets/ds001021/versions/1.0.0

https://nilab-uva.github.io/AOMIC.github.io/

Data of the 1000Brains are available upon request from the responsible Principal Investigator (Caspers et al., 2014).

The code for preprocessing, data preparation, model training and computation of further analyses is available on Github:

https://jugit.fz-juelich.de/l.wiersch/functional_sex_classification_code

https://jugit.fz-juelich.de/f.hoffstaedter/bids_pipelines/-/tree/master/func

