## Supplementary Material for "Sex classification from functional brain connectivity: Generalization to multiple datasets"

This file includes:

Figure S1 to S3
Table S1 to S2

**Supplementary Results**

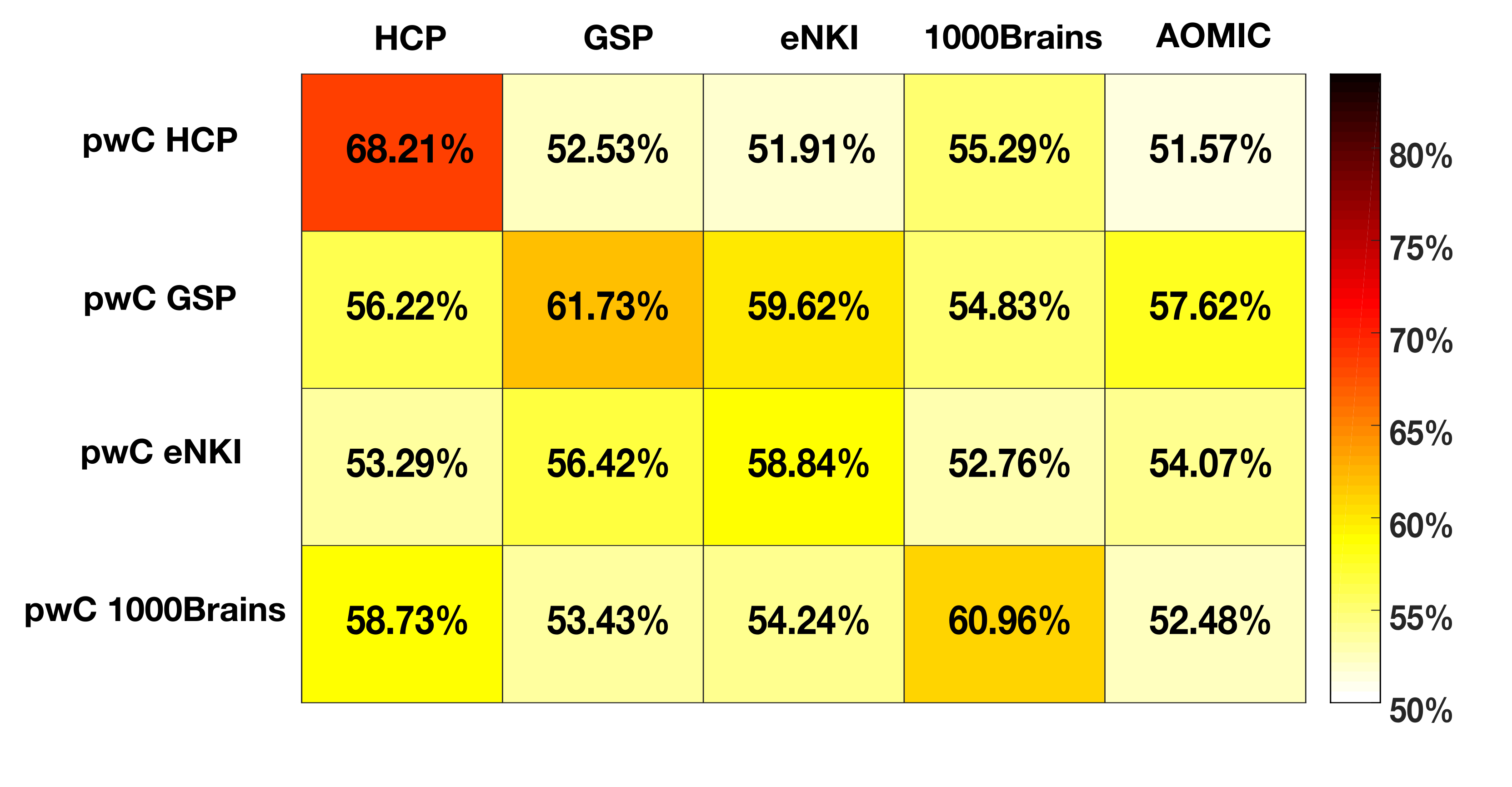

**Figure S1. Tile Plots of mean accuracies.** Mean accuracies averaged across all 436 parcels for each CV- and across-sample predictions of pwCs trained on the data of single samples.

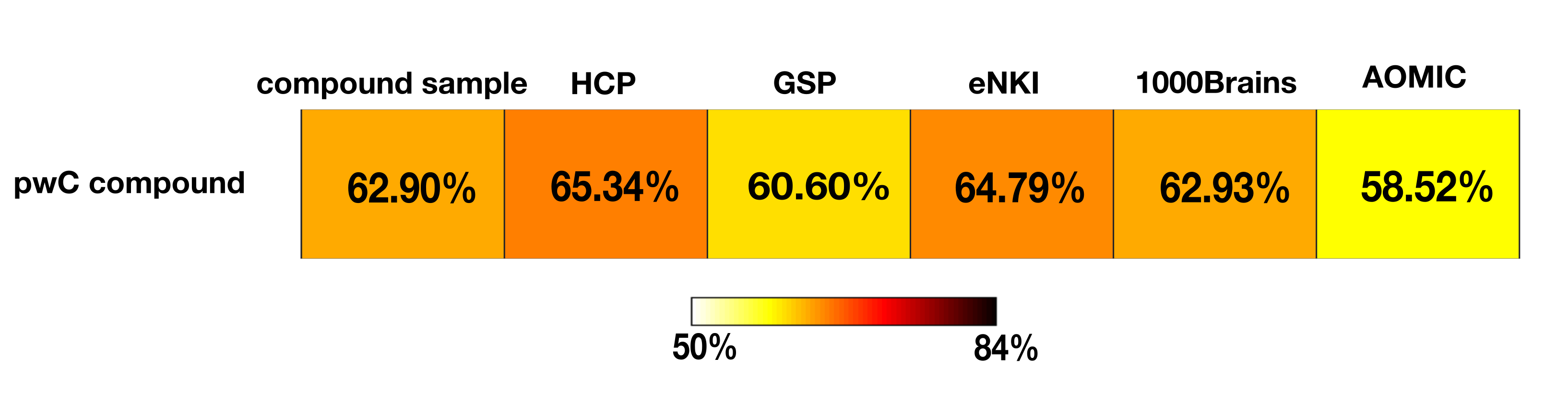

**Figure S2. Tile Plots of mean accuracies.** Mean accuracies averaged across all 436 parcels for the CV- and across-sample predictions of pwC compound.

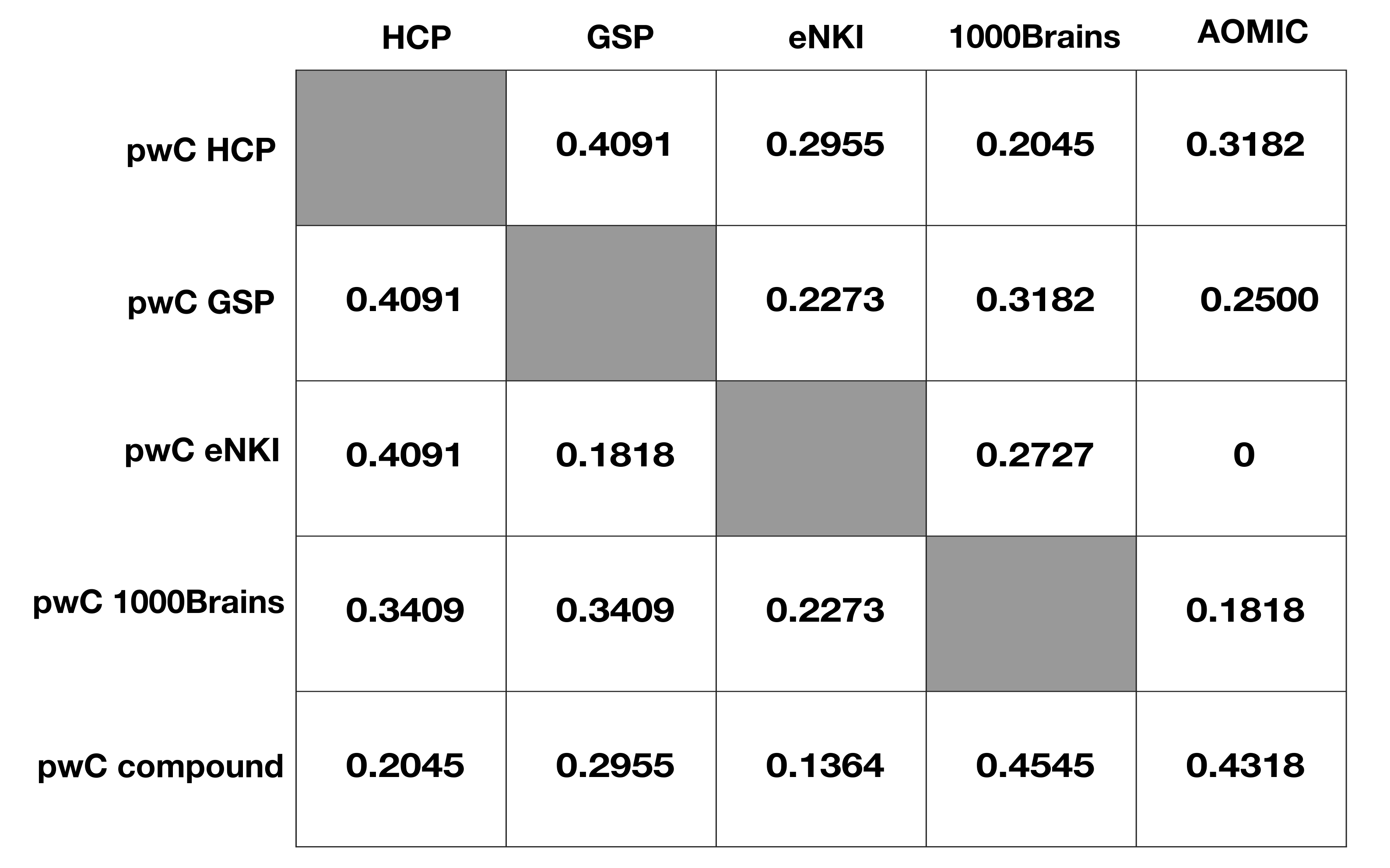

**Figure S3. Spatial consistency of all pwCs.** For each combination of training (rows) and test sample (columns), dice coefficients were calculated as the number of similar parcels classifying within the top 10% parcels for each CV- and across-sample prediction.

| **Table S1** |  |  |  |  |  |  |  |  |  |  |  |  |
| --- | --- | --- | --- | --- | --- | --- | --- | --- | --- | --- | --- | --- |
| Mean and range in percent of sex classification accuracies for within- and between-dataset predictions trained on single datasets (model 1-4) | | | | | | | | | | | | |
|  |  |  |  | **Model applied to** | | | | | | |  |  |
|  |  | **compound sample** |  | **HCP** |  | **GSP** |  | **eNKI** |  | **1000Brains** |  | **AOMIC** |
| **pwC Map HCP** |  |  |  | 68.2 (51.0 - 77.4) |  | 52.5 (49.0 - 64.0) |  | 51.9 (44.7 - 65.3) |  | 55.3 (49.0 - 61.8) |  | 51.6 (47.1 - 65.0) |
| **pwC Map GSP** |  |  |  | 56.2 (46.0 - 67.3) |  | 61.7 (52.4 - 68.5) |  | 59.6 (49.0 - 70.0) |  | 54.8 (48.1 - 64.1) |  | 57.6 (50.0 - 68.1) |
| **pwC Map eNKI** |  |  |  | 53.3 (46.0 - 63.9) |  | 56.4 (48.7 - 63.8) |  | 58.8 (47.3 - 72.1) |  | 52.8 (45.8 - 62.5) |  | 54.1 (45.2 - 65.0) |
| **pwC Map 1000Brains** |  |  |  | 58.7 (49.4 - 68.7) |  | 53.4 (49.5 - 65.2) |  | 54.2 (47.4 - 66.8) |  | 61.0 (54.2 - 69.3) |  | 52.5 (46.5 - 66.2) |
| **pwC Map compound** |  | 62.9 (53.7 - 70.1) |  | 65.3 (45.9 - 74.6) |  | 60.6 (49.1 - 70.1) |  | 64.8 (47.9 - 83.3) |  | 62.9 (52.4 - 75.2) |  | 58.5 (49.3 - 69.0) |

| **Table S2.** Comparisons of performance of the 10% best classifying parcels each pwC on the different test samples (a) and comparisons of performance of the different pwCs on each test samples (b) | | | | |
| --- | --- | --- | --- | --- |
| **(a)** |  |  |  |  |
| **pwC HCP** |  |  |  |  |
|  | **AOMIC** | **1000Brains** | **eNKI** |  |
| **GSP** | t = 1.40, p = 0.1643 | t = -2.23, p = 0.0281 | t = 1.17, p = 0.2464 |  |
| **eNKI** | t = 0.33, p = 0.7387 | t = -3.58, p = 0.0005 |  |  |
| **1000Brains** | t = 3.50, p = 0.0007 |  |  |  |
| **pwC GSP** |  |  |  |  |
|  | **AOMIC** | **1000Brains** | **eNKI** |  |
| **HCP** | t = 1.87, p = 0.0649 | t = 9.97, p < 0.0001 | t = -7.79, p < 0.0001 |  |
| **eNKI** | t = 9.27, p < 0.0001 | t = 16.81, p < 0.0001 |  |  |
| **1000Brains** | t = -7.83, p < 0.0001 |  |  |  |
| **pwC eNKI** |  |  |  |  |
|  | **AOMIC** | **1000Brains** | **GSP** |  |
| **HCP** | t = 2.88, p = 0.0050 | t = 5.48, p < 0.0001 | t = -1.83, p = 0.0711 |  |
| **GSP** | t = 5.66, p < 0.0001 | t = 8.76, p < 0.0001 |  |  |
| **1000Brains** | t = -2.96, p = 0.0039 |  |  |  |
| **pwC 1000Brains** |  |  |  |  |
|  | **AOMIC** | **eNKI** | **GSP** |  |
| **HCP** | t = 13.02, p < 0.0001 | t = 6.71, p < 0.0001 | t = 14.03, p < 0.0001 |  |
| **GSP** | t = 1.67, p = 0.0992 | t = -5.85, p < 0.0001 |  |  |
| **eNKI** | t = 6.56, p < 0.0001 |  |  |  |
|  | alpha level: 0.008 |  |  |  |
| **pwC compound** |  |  |  |  |
|  | **AOMIC** | **1000Brains** | **eNKI** | **GSP** |
| **HCP** | t = 21.05, p < 0.0001 | t = 6.55, p < 0.0001 | t = -6.11, p < 0.0001 | t = 16.74, p < 0.0001 |
| **GSP** | t = 5.25, p < 0.0001 | t = -8.64, p < 0.0001 | t = -17.29, p < 0.0001 | |
| **eNKI** | t = 20.42, p < 0.0001 | t = 10.46, p < 0.0001 |  |  |
| **1000Brains** | t = 13.03, p < 0.0001 |  |  |  |
|  | alpha level: 0.005 |  |  |  |

**(b)**

| **test sample HCP** |  |  |  |  |
| --- | --- | --- | --- | --- |
|  | **pwC compound** | **pwC 1000Brains** | **pwC eNKI** |  |
| **pwC GSP** | t = -29.57, p < 0.0001 | t = -4.54, p < 0.0001 | t = 11.23, p < 0.0001 |  |
| **pwC eNKI** | t = -36.27, p < 0.0001 | t = -15.18, p < 0.0001 |  |  |
| **pwC 1000Brains** | t = 25.64, p < 0.0001 |  |  |  |
| **test sample GSP** |  |  |  |  |
|  | **pwC compound** | **pwC 1000Brains** | **pwC eNKI** |  |
| **pwC HCP** | t = -22.16, p < 0.0001 | t = -3.68, p = 0.0004 | t = -4.84, p < 0.0001 |  |
| **pwC eNKI** | t = -24.64, p < 0.0001 | t = 0.71, p = 0.4816 |  |  |
| **pwC 1000Brains** | t = -20.43, p < 0.0001 |  |  |  |
| **test sample eNKI** |  |  |  |  |
|  | **pwC compound** | **pwC 1000Brains** | **pwC GSP** |  |
| **pwC HCP** | t = -31.93, p < 0.0001 | t = -9.77, p < 0.0001 | t = -19.01, p < 0.0001 |  |
| **pwC GSP** | t = -19.97, p < 0.0001 | t = 9.33, p < 0.0001 |  |  |
| **pwC 1000Brains** | t = -25.36, p < 0.0001 |  |  |  |
| **test sample 1000BRrains** | |  |  |  |
|  | **pwC compound** | **pwC eNKI** | **pwC GSP** |  |
| **pwC HCP** | t = -35.92, p < 0.0001 | t = 5.66, p < 0.0001 | t = -4.22, p < 0.0001 |  |
| **pwC GSP** | t = -27.70, p < 0.0001 | t = 8.25, p < 0.0001 |  |  |
| **pwC eNKI** | t = -35.29, p < 0.0001 |  |  |  |
|  | alpha level: 0.008 |  |  |  |
| **test sample AOMIC** |  |  |  |  |
|  | **pwC compound** | **pwC 1000Brains** | **pwC eNKI** | **pwC GSP** |
| **pwC HCP** | t = -16.08, p < 0.0001 | t = -2.75, p = 0.0072 | t = -2.01, p = 0.0476 | t = -11.02, p < 0.0001 |
| **pwC GSP** | t = -7.78, p < 0.0001 | t = 8.41, p < 0.0001 | t = 13.60, p < 0.0001 |  |
| **pwC eNKI** | t = -20.71, p < 0.0001 | t = -1.37, p = 0.1745 |  |  |
| **pwC 1000Brains** | t = -13.92, p < 0.0001 |  |  |  |
|  | alpha level: 0.005 |  |  |  |
